# KGETCDA: an efficient representation learning framework based on knowledge graph encoder from transformer for predicting circRNA-disease associations

**DOI:** 10.1101/2023.03.28.534642

**Authors:** Jinyang Wu, Zhiwei Ning, Yidong Ding, Ying Wang, Qinke Peng, Laiyi Fu

## Abstract

Recent studies have demonstrated the significant role that circRNA plays in the progression of human diseases. Identifying circRNA-disease associations (CDA) in an efficient manner can offer crucial insights into disease diagnosis. While traditional biological experiments can be time-consuming and labor-intensive, computational methods have emerged as a viable alternative in recent years. However, these methods are often limited by data sparsity and their inability to explore high-order information. In this paper, we introduce a novel method named Knowledge Graph Encoder from Transformer for predicting CDA (KGETCDA). Specifically, KGETCDA first integrates more than 10 databases to construct a large heterogeneous non-coding RNA dataset, which contains multiple relationships between circRNA, miRNA, lncRNA and disease. Then, a biological knowledge graph is created based on this dataset and Transformer-based knowledge representation learning and attentive propagation layers are applied to obtain high-quality embeddings with accurately captured high-order interaction information. Finally, multilayer perceptron is utilized to predict the matching scores of CDA based on their embeddings. Our empirical results demonstrate that KGETCDA significantly outperforms other state-of-the-art models. To enhance user experience, we have developed an interactive web-based platform named HNRBase that allows users to visualize, download data and make predictions using KGETCDA with ease.

## INTRODUCTION

Circular RNA (circRNA) is a unique type of endogenous non-coding RNA (ncRNA) molecules typically characterized by a covalently closed-loop structure without 3’ and 5’ polyadenylated tails (1). With the rapid development of high-throughput sequencing technologies, increasing studies have demonstrated that circRNA is extensively involved in numerous physiological and pathological processes in humans, including regulating gene expression (2), acting as miRNA sponges (3) and mediating cancer metastasis (4). Thus, exploring the associations between circRNAs and diseases can offer crucial insights into disease pathology at molecular resolution and help identify promising biomarkers for clinical diagnosis and treatment of complex diseases (5). However, despite the extensive cataloging of circRNAs (6, 7), the number of experimentally confirmed disease-related circRNAs (8, 9, 10) remains relatively small due to the time-consuming and expensive laboratory experiments.

To solve this problem, a series of computational methods have been proposed to identify potential circRNA-disease associations (CDA) recently. These approaches could be roughly divided into three types, including network-based, traditional machine learning-based and deep learning-based methods. Firstly, network-based algorithms construct heterogeneous networks based on known CDA and biological similarity calculation to predict CDA. For example, KATZHCDA (11), KATZCPDA (12), and IBNPKATZ (13) employ the KATZ algorithm to infer potential CDA. Random walk and label propagation are other frequently used methods. Huseyin et al. (14) applied random walk with restart (RWR) to predict CDA based on similarity matrices. Zhang et al. (15) adopted a linear neighborhood label propagation method (CD-LNLP) to identify potential CDA after constructing similarity networks. Secondly, traditional machine learning-based algorithms identify CDA by training classifiers after handcrafted feature construction. Lei et al. (16) proposed GBDTCDA to predict CDA, where gradient boosting decision tree was employed on manually constructed fusion features. Zhang et al. (17) used the weighted nuclear norm minimization to identify associations between circRNAs and diseases. Moreover, Matrix factorization (MF) have also been proposed for CDA identification recently due to MF’s exceptional performance in exploring latent information from sparse and noisy data. Wei et al. (18) designed a new computational method (iCircDA-MF) to calculate affinity scores of CDA using MF. Peng et al. (19) combined robust nonnegative matrix factorization with label propagation (RNMFLP) to infer potential CDA. Thirdly, deep-learning based algorithms have demonstrated the superiority in automatically extracting latent features in many fields (20, 21, 22), which is also a significant branch in the domain of CDA prediction (23, 24, 25, 26). Specially, Lu et al. (24) put forward a computational model (CDASOR) based on convolutional and recurrent neural networks and Deepthi et al. (25) applied deep autoencoder to identify CDA (AE-RF). Considering the graph structure consisting of CDA, Niu et al. (27) used the graph markov neural network (GMNN) to capture deep representations of CDA and make predictions. Lan et al. (28) designed the knowledge graph with attention mechanism (KGANCDA) for inferring unknown CDA.

Although existing methods have achieved significant progress in the field of CDA prediction, they still face limitations due to data sparsity and their inability to efficiently explore high-order information that combines biological data characteristics. For example, most existing methods rely heavily on similar circRNAs to predict disease-related circRNAs, which can be challenging when working with sparse data and unable to effectively explore high-order information. Therefore, it is necessary to incorporate other circRNA-related relations and construct a large heterogeneous dataset to alleviate data sparsity and improve performance of current models to some extent. Additionally, it’s also necessary and significant to propose a method capable of efficiently processing multi-source data and extracting high-order information by integrating the intrinsic properties of biological data.

In this paper, we introduce a new framework for predicting CDA called Knowledge Graph Encoder from Transformer for CDA (KGETCDA). Our framework is based on representation learning and can effectively capture high-order interaction information between multi-source data and the intrinsic properties of biological molecules. We start by integrating over 10 databases to construct a heterogeneous ncRNA dataset, including associations between circRNA, miRNA, lncRNA and disease. After that, a biological knowledge graph is created based on the dataset. Additionally, knowledge graph representation learning and attentive embedding propagation layers are designed to generate optimal embeddings with captured high-order connectivity. Finally, multilayer perceptron is applied to obtain reliable matching scores of CDA based on their embeddings. The empirical experiments demonstrate that KGETCDA outperforms other state-of-the-art (SOTA) models. Moreover, we develop a user-friendly web-based platform for users to visualize, download data and make predictions using KGETCDA with ease. The key contributions of our work are summarized as follows:

- We integrate multi-source data (more than 10 databases) on circRNA, miRNA, lncRNA and disease to construct a large heterogeneous ncRNA dataset, which helps alleviate data sparsity and benefits future research in this field (previous methods use less than five databases).
- Our proposed method KGETCDA is based on a transformer-based knowledge graph representation learning framework that effectively captures high-order information and intrinsic properties of biological data.
- Extensive experiments and case studies are conducted on two datasets, demonstrating the effectiveness of KGETCDA and its potential to assist future research on circRNA-miRNA-lncRNA axes in pathogenesis.
- We develop an interactive web-based platform for users to visualize, download data and make predictions using KGETCDA conveniently, which enhances user experience and promote diversity in presenting results in this filed.
- Our proposed framework’s strong performance can also encourage the development of other biological fields, and we have investigated its use in the field of miRNA-disease and lncRNA-disease prediction.

The remaining part of this paper proceeds as follows. The section ‘MATERIALS AND METHODS’ introduces the constructions of datasets, similarity measures and the overall architecture of KGETCDA. Then, evaluation criteria, performance comparison with SOTA methods, ablation experiments, parameter setting and case studies are discussed in the section ‘RESULTS’. After that, we present an interactive web-based platform for users in ‘WEB-BASED VISUALIZATION’. Finally, some discussions on KGETCDA and future works are introduced in ‘DISCUSSION AND CONCLUSION’.

## MATERIALS AND METHODS

In this section, we present the proposed method KGETCDA, which efficiently captures high-order interaction information. Figure 1 illustrates the model framework, consisting of three parts: data collection, similarity measures and model prediction. Firstly, we collect multi-source data and construct a large, heterogeneous ncRNA dataset. Next, we prepare similarity measures for negative sampling. Subsequently, a biological knowledge graph is created, and optimal embeddings are obtained using Transformer-based knowledge representation learning and attentive embedding propagation layers. Finally, MLP-based prediction layers are applied to output the affinity scores of candidate CDA.

**Figure 1.**
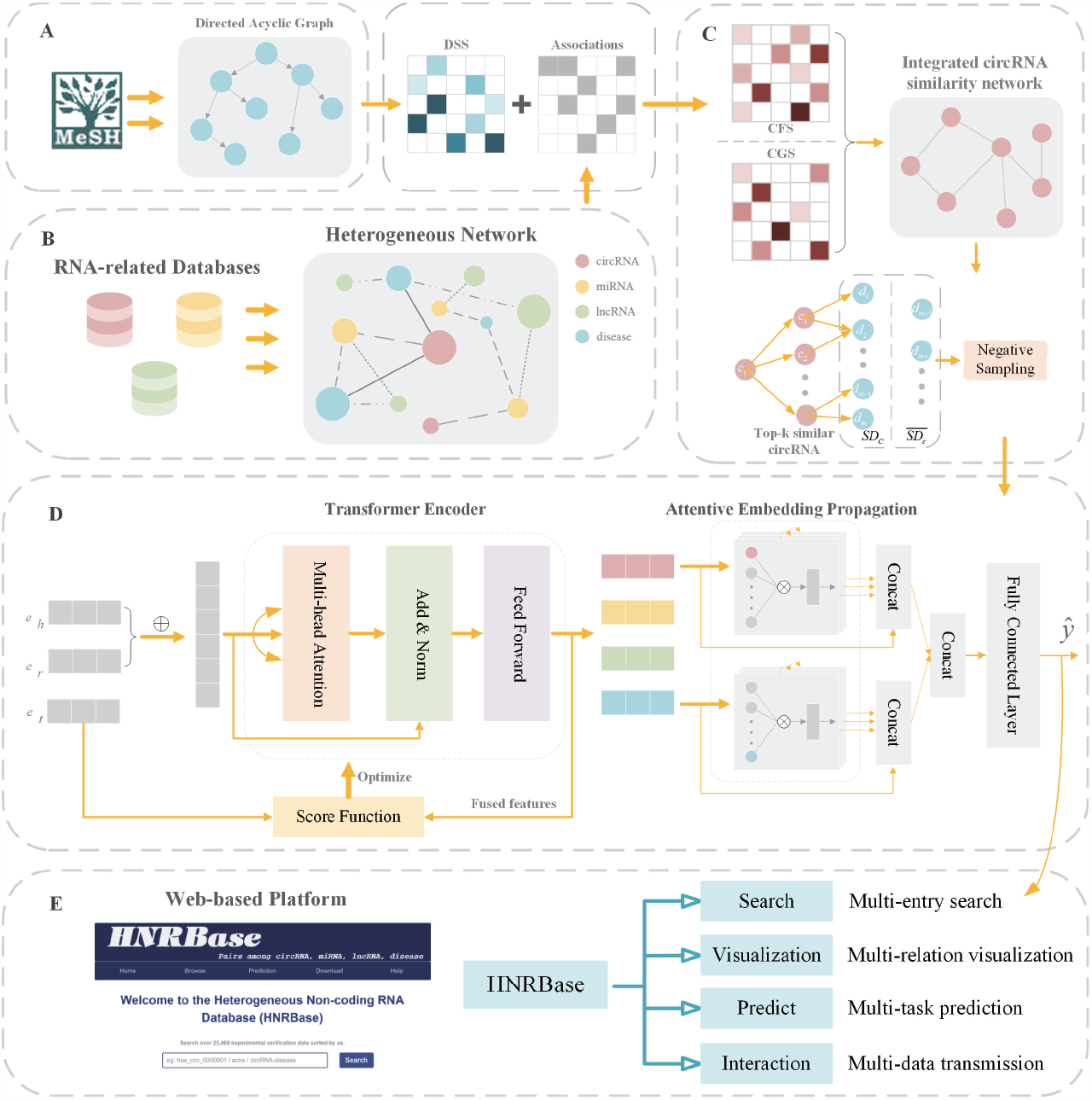
The flowchart of KGETCDA. **(A)** Extract disease information by creating a directed acyclic graph. **(B)** Construct a heterogeneous network via integrating multi-source ncRNA-disease databases. **(C)** Similarity-based negative sampling. It first constructs the integrated similarity descriptor, and then obtains top-k similar circRNAs of candidate circRNA *i*. Finally, negative samples (diseases) are selected from unknown associated diseases of these diseases. **(D)** Produce high-quality embeddings and identify potential circRNA-disease associations. **(E)** Web-based Visualization.

### Datasets

In this study, we first obtain 330 circRNAs, 265 miRNAs, 404 lncRNAs, 79 diseases, 346 circRNA-diseases associations, 106 miRNA-disease associations, 527 lncRNA-disease associations, 146 circRNA-miRNA associations and 202 miRNA-lncRNA associations from previous work (28) to get a commonly used dataset in previous works for dataset1. Then, for dataset2, we integrate multi-source data from 13 databases to construct a larger heterogeneous dataset. Specifically, for dataset2, we gather associations of circRNA-disease and circRNA-miRNA from Circad (10), MNDR (29), Lnc2Cnacer (30), LncRNADisease (31), CircRNADisease (32), Circ2Disease (8) and CircR2Cancer (9). And the associations of miRNA-disease, miRNA-lncRNA, lncRNA-disease are obtained from MNDR (29), HMDD (33), Circ2Disease (8), StarBase (34), LncRNADisease (31) and Lnc2Cancer (30). We ensure the consistency by matching circRNA names with circBase (6) and circBank (7) as much as possible, addressing the inconsistent standard issue of current circRNA-disease datasets. Additionally, most of the disease names in dataset1 have been replaced with those in disease ontology (35). After removing redundant associations, 561 circRNAs, 658 miRNAs, 1043 lncRNAs, 190 diseases,

1399 circRNA-disease associations, 10154 miRNA-disease associations, 3280 lncRNA-disease associations, 1129 circRNA-miRNA associations and 9506 miRNA-lncRNA associations are obtained for dataset1. Table 1 summarizes the information of the two datasets. To visualize datasets, we show the degree distribution of diseases and the relationship between circRNA-disease and circRNA-position of dataset1 in Figure 2A, 2B and dataset2 in Supplementary Figure S1, and the process of constructing the dataset2 is shown in Figure 2C. We observe that some diseases associate with more circRNAs than others, highlighting the significance of designing an effective architecture capable of extracting features from the above complex multi-relational scene.

**Table 1.** The summary information of two datasets

| Dataset | circRNA-disease | miRNA-disease | lncRNA-disease | circRNA-miRNA | miRNA-lncRNA | total |
| --- | --- | --- | --- | --- | --- | --- |
| Dataset1 | 346 | 106 | 527 | 146 | 202 | 1327 |
| Dataset2 | 1399 | 10154 | 3280 | 1129 | 9506 | 25468 |

**Figure 2.**
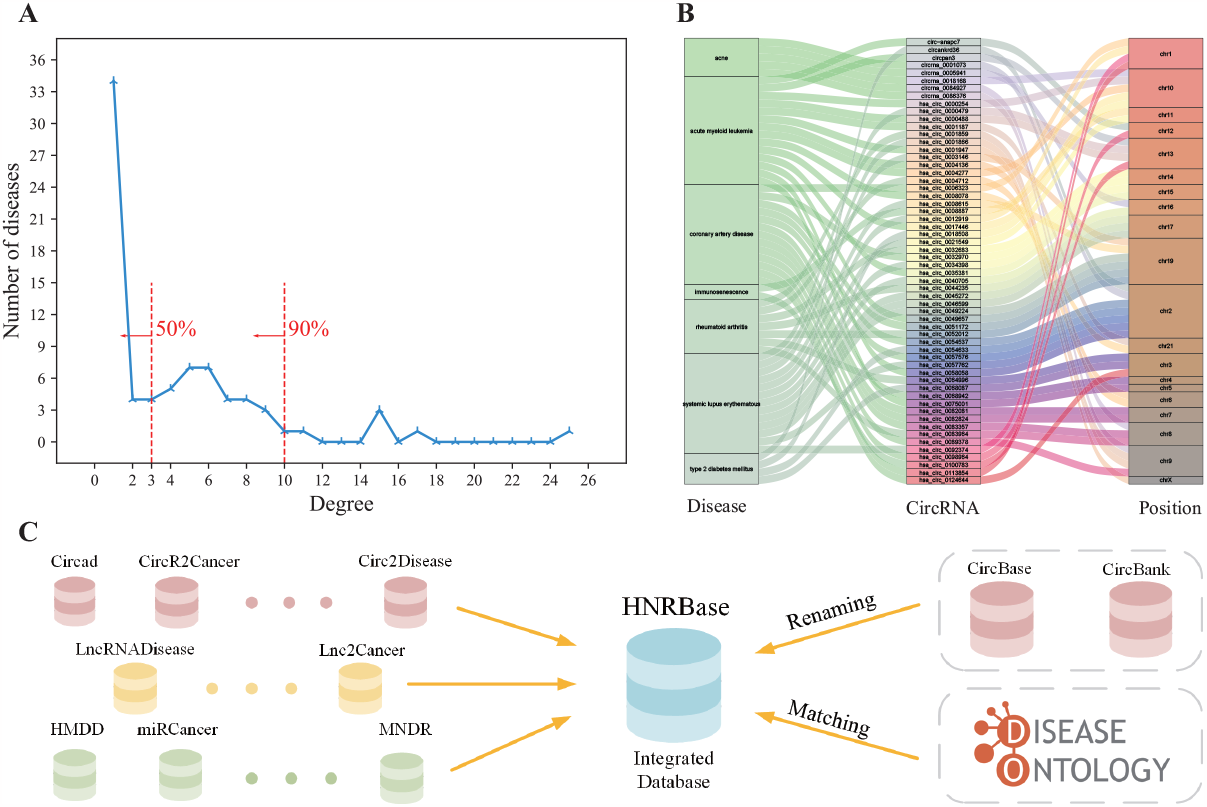
Visualization of dataset1. **(A)** The degree distribution. It represents the number of corresponding diseases associating with different numbers of circRNAs in dataset1, where fifty percent of diseases only associate with less than 3 circRNAs and ninety percent of diseases associate with up to 10 circRNAs. **(B)** The visualization of sankey diagram of some associations in dataset1. **(C)** The process of constructing dataset2.

### Similarity measures

To obtain intrinsic properties of multi-source biological data, an integrated feature descriptor is constructed based on disease semantic similarity, circRNA functional similarity and gaussian interaction profile kernel similarity.

#### Disease semantic similarity

Following previous studies (25, 36, 37), the disease information is obtained by calculating disease semantic similarity based on the medical subject headings (MeSH), which is provided by the National Library of Medicine (https://www.ncbi.nlm.nih.gov/). In the MeSH database, diseases are described as a directed acyclic graph (DAG), where nodes represent diseases and edges represent relationships between diseases. For example, given a disease *d*, it can be represented as *DAG*_*d*_ =(*d,A*_*d*_,*R*_*d*_), where *A*_*d*_ represents the ancestral node set of *d* including itself and *R*_*d*_ represents the set of *d*’s corresponding relationships. Additionally, there are two approaches to calculate disease semantic similarity from different perspectives.

The first method is based on the assumption that different ancestral nodes of disease *d* in the same layer have the same contribution to *d*. Specially, assuming that another disease *k* exits in *DAG*_*d*_, its contribution to disease *d* is described by:

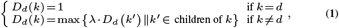

where *λ* controls the semantic contribution between disease *k* and its child diseases. Similar to previous work (36), *λ* is set to 0.5. Then, we can obtain the semantic value of disease *d* as follows:

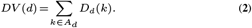

Finally, for disease *m* and *n*, we can calculate their semantic similarity based on the assumption that two diseases sharing larger parts in DAG are more similar. The formula is

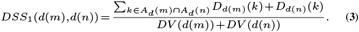

The second method considers that diseases appearing less in DAG may be more important. Therefore, the contribution of disease *k* can be calculated as follows:

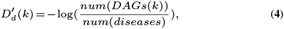

where *num*(*DAGs*(*k*)) denotes the number of *DAGs* containing disease *k*, and *num*(*diseases*) denotes the number of all diseases. Then, similar to the first method, the second semantic similarity can be calculated by

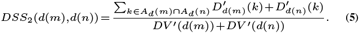

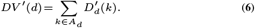

#### CircRNA functional similarity

Under the hypothesis that circRNAs sharing similar disease groups tend to be functionally similar (36), we can obtain the functional similarity matrix for circRNA. Specifically, we define *D*_*i*_ and *D*_*j*_ to represent the disease set related to circRNA *c*_*i*_ and *c*_*j*_, respectively. Then, the functional similarity between circRNA and can be described as follows:

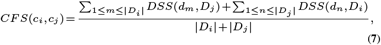

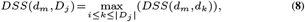

where *DSS*(*d*_*m*_,*D*_*j*_) is the semantic similarity value between disease *d*_*m*_ and disease group *D*_*j*_.

#### Gaussian interaction profile kernel similarity

Assuming that similar circRNAs associate with similar diseases and vice versa, the gaussian interaction profile kernel similarity (GIPKS) for circRNAs can be calculated (38). Here, we define the association matrix 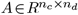, in which *n*_*c*_ and *n*_*d*_ denote the total number of circRNAs and diseases, respectively. Then the GIPKS for circRNA *c*_*i*_ and *c*_*j*_ can be obtained by

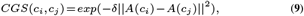

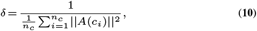

where *n*_*c*_ is the number of all circRNAs, *δ* controls the kernel bandwidth, and *A*(*c*_*i*_) and *A*(*c*_*j*_) denote the *i*th and *j*th rows of *A*, respectively.

#### Similarity fusion

To obtain a reasonable representation of disease semantic similarity, element-wise averaging of the above two disease similarity methods is applied to construct the disease semantic similarity descriptor:

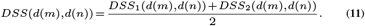

Then, considering the information of different similarity methods in different feature spaces, similarities above are integrated to produce the fusion feature descriptor for circRNAs:

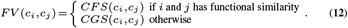

### KGETCDA

We now introduce the method KGETCDA and its detailed implementation here, which captures high-order interaction information in an end-to-end manner. An overview of the KGETCDA architecture is presented in Figure 1D, which consists of four parts: knowledge graph encoder from Transformer, attentive information propagation, MLP-based association prediction and negative sampling based on similarity measuring.

#### Knowledge graph encoder from Transformer

Recently, knowledge graph (KG) as a form of structured human knowledge has been widely applied to various applications such as question answering (39), recommendation (40) and time series prediction (41). Due to KG’s outstanding feature representation capability, they are also gradually applied to computational biology like drug-drug prediction (42) and drug-target protein prediction (43). For our research, we can create a heterogeneous KG based on datasets construct before (illustrated in Figure 3). Specially, the KG of ncRNA-disease can be described as *G* =(*E,R,F*), where *E* denotes the entity set (circRNA, miRNA, lncRNA, disease), *R* denotes the relation set (associations between entities), and *F* denotes the fact set (a fact is a triplet (*h,r,t*) ∈ *F,h* ∈ *E,r* ∈ *R,t* ∈ *E*).

**Figure 3.**
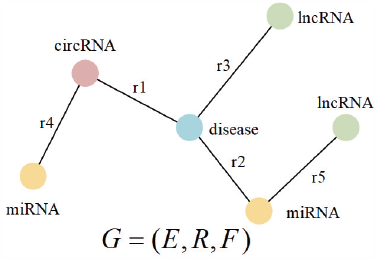
The overview of knowledge graph. It contains 4 entities (circRNA, miRNA, lncRNA, disease) and 5 relations (circRNA-disease, miRNA-disease, lncRNA-disease, circRNA-miRNA, miRNA-lncRNA).

Acknowledged as the pivotal research issue of KG, knowledge representation learning (KRL, also known as knowledge graph embedding) is an effective tool to model complex relational graphs like biological systems and produce reliable representations of entities and relations for downstream tasks. Furthermore, due to Transformer’s (44) remarkable performance in computer vision (45) and natural language processing (21, 22), considerable efforts have been devoted to integrating Transformer into KRL (46, 47). Inspired by these excellent works, we propose a Transformer encoder-based architecture. As illustrated in Figure 1D, multi-head self-attention mechanism of Transformer encoders is applied to effectively encode KG and produce high-quality embeddings. In particular, for a given triple 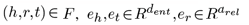 are defined as the embedding vector for head entity, tail entity and relation, respectively. For simplicity, we assume that *d*_*ent*_ = *d*_*rel*_ = *d*. Then, the concatenated embeddings *CE* ∈ *R*^2*×d*^ of the pair (h,r) are considered to be the input sequence, which also serves as the query *Q*, the key *K* and the value *V* for self-attention mechanism. Therefore, in order to obtain highly expressive information from different representation subspaces, multi-head attention mechanism is used as follows:

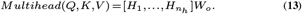

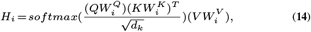

where [] represents the concatenation operation, *n*_*h*_ represents the number of heads, and 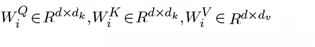 and 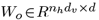 are linear projection matrices. Additionally, more attention heads are applied than the standard Transformer (44), since increasing the number of attention heads would offer better representations and facilitate latent feature extraction in different subspaces (46). Afterwards, a token-wise feed-forward network following is utilized to further enhance model’s capability to produce powerful representations and perform various complex tasks.

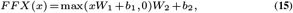

where *x* is the output of multi-head attention mechanism, and 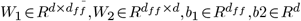 are parameter matrices. Note that Yun et al. (48) demonstrated Transformer’s universal approximation power in various sequence-based tasks with the architecture of multi-head attention mechanism combined with a fully connected feed-forward network.

After obtaining learned embeddings of entities and relations based on the above architecture, a score function is designed for measuring the plausibility of a given triple (*h,r,t*) similar to previous work (49, 51). In this paper, we use a simple method named TwoMult (46, 50), which is formulated as follows:

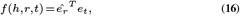

where 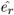 is the updated embedding of the relation passing the Transformer-encoder based architecture above and *e*_*t*_ is the initial embedding of tail entity. Despite its simplicity, TwoMult is an effective method to decode embeddings and produce reliable matching scores of (*h,r,t*) (46). Moreover, it’s our belief that the Transformer encoder-based architecture has perfectly embedded the information of head entity into the relation embedding and thus we only use the updated relation embedding in TwoMult. Noteworthy, we use the simple inner product as scoring function here for simplicity, and leave further exploration as future works. Finally, a margin-based ranking criterion (49) following is applied to train KRL part, which aims to encourage the discrimination between positive and negative triples.

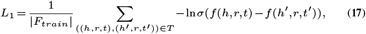

where *σ* is the sigmoid function, |*F*_*train*_| is the number of all facts in training set, and *T* is the set of all positive triples and equal proportion of negative triples, which are constructed by replacing head or tail entity in a positive triple once. A positive triple will be assigned a higher score while a negative triple gets a lower score. In summary, the architecture of knowledge graph encoder from Transformer constructs a hierarchical representation of complex biological KG by effectively merging high-order neighborhood information, which paves the way for downstream tasks such as CDA prediction.

#### Attentive information propagation

Considering that our downstream task is CDA prediction and the properties of each entity in the above KG structure have no difference by default, we build an attentive information propagation architecture following (40) to further distinguish different neighbors and strengthen the representation of two specific entities, i.e. circRNA and disease. In the following, we will start with a single layer, consisting of two parts: information propagation and information aggregation, and then extend to multiple layers.

### Information propagation

To obtain high-quality expression, it’s crucial to fully exploit the information of neighbors of an entity. Therefore, we will describe how to generate an efficient combination of neighborhood information in this part. Specifically, given an entity *h* (circRNA or disease), we can define its ego-network *N*_*h*_ = { (*h,r,t*) |*h,t* ∈ *E,r* ∈ *R*,(*h,r,t*) ∈ *F*} (52), which represents triples consisting of the entity *h*, all neighbors of *h* and their relations. Then, the integrated information of *h*’s ego-network can be defined as follows:

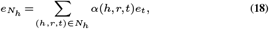

where *e*_*t*_ is the embedding of tail entity defined before, and *α*(*h,r,t*) is the propagation score deciding how much information is propagated from neighbor *t* to *h*. Since different neighbors have different importance (40, 54), knowledge-aware attention is applied to calculate the propagation score, which is formulated as follows:

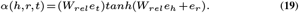

where *W*_*rel*_ is the transformation matrix, which projects entity embeddings into the relation space. With the relational attention mechanism (40), the process of masked information propagation becomes efficient. Moreover, normalization by using the following softmax function is applied on propagation scores to reinforce their asymmetry and thus distinguish the importance of different neighbors to capture effective information.

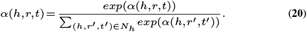

### Information aggregation

After the information propagation, we can aggregate the representation of the entity *h* and its ego-network. Graph convolutional network (GCN) aggregator (55) is employed, which is defined as follows:

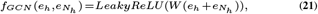

where 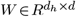 is the trainable matrix and *d*_*h*_ is the transformation size. Moreover, GraphSage aggregator (53) and Bi-Interaction aggregator (40) following are applied to compare with GCN aggregator to demonstrate the effectiveness of the GCN aggregator in our research.

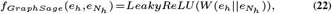

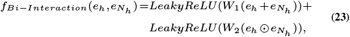

where || denotes the concatenation operation and ⊙ denotes the element-wise product.

After obtaining the first-order interaction information, we can stack more layers to further strengthen high-order representations of circRNAs and diseases, which contain abundant information from their multi-hop neighbors. In particular, for the entity *h*, the representation of *l*-th layer can be described as follows:

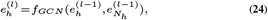

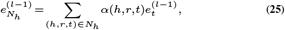

where 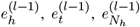 is the embedding of entity *h, t*, and ego-network of *h* in the (*l* − 1)-th layer, respectively. Clearly, after performing above attentive propagation layer architecture, the high-order interaction information is further explored, and the encoding representation of circRNAs and diseases are more expressive and reliable. Following (28, 40), *l* is set to 4 because the fourth-order information is sufficient for accurately exploiting latent features in KG.

#### MLP-based association prediction

With above multiple attentive propagation layers, high-quality embeddings of each entity (circRNA and disease) with accurately captured high-order information are obtained. Note that the embeddings of different layers represent the interaction information of corresponding orders, the concatenation operation of these embeddings is applied to further enrich the final representation and adjust the strength of information propagation as follows:

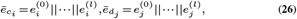

where || denotes the concatenation operation. Similar to (23, 25, 28), we also concatenate the embedding 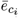 and 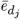 of the two entity to get fusion descriptor of a candidate CDA. Then, the multilayer perceptron is applied on the concatenated features to produce reliable affinity scores, which is formulated as follows:

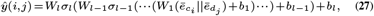

where *W*_*l*_, *b*_*l*_ and *σ*_*l*_ are the trainable parameter matrix, the bias, and the activation function in the *l*-th linear layer. Moreover, we select ReLU (56) as the activation function in hidden layers and sigmoid in the output layer to control the output to be a probability value from 0 to 1. To optimize this part, we get the BPR loss (57) as:

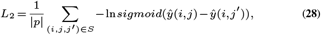

where |*P*| is the number of all pairs consisting of head entities and tail entities in training set, and *S* is the set of all positive pairs and negative triples of equal proportion, which represents unobserved triples here.

Finally, to achieve the joint model optimization in an end-to-end manner, the whole loss is defined as follows:

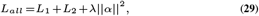

where *λ* is a regularization parameter to prevent overfitting, *α* is all parameters of the model, and *L*_1_ and *L*_2_ are the margin-based ranking loss and BPR loss, respectively.

#### Negative Sampling based on similarity measuring

Recent studies have demonstrated the significant role that negative sampling plays in efficient model learning in various fields, including knowledge graph embedding (49), recommendation (59) and computational biology (26, 58). High-quality negative sampling can effectively facilitate the model learning towards correct direction. However, exiting methods commonly rely on a random negative sampling strategy, which selects unobserved CDA as negative samples randomly and thus often result in unidentified false negative signals. Therefore, following (26), a similarity-based negative sampling strategy is applied in this paper. Specifically, given that similar circRNAs tend to associate with similar diseases and vice versa, we can obtain integrated similarity descriptor in the part ‘Similarity measures’. Afterwards, considering a circRNA *i*, we use *SC* = {*c*_1_,*c*_2_, ·,*c*_*k−*1_,*c*_*k*_} and *SD* = {*d*_1_,*d*_2_, ·,*d*_*n*_} to denote the set of the top-k most similar circRNAs selected based on the similarity descriptor, and the set of all diseases, respectively. Then, the related diseases *SD*_*c*_ = {*d*_1_,*d*_2_·,*d*_*m*_} of all circRNAs in *SC* in the training set can be obtained and they will not be chosen to consist of the negative CDA. We generate a negative pair (*c*_*i*_,*d*_*j*_) only by selecting the disease *j* from *SD* − *SD*_*c*_ = {*d*_*m*+1_,*d*_*m*+2_, ·,*d*_*n*_}. The specific architecture of negative sampling is illustrated in Figure 1C. It’s our belief that this similarity-based strategy could produce high-quality negative samples for better model learning and thus effectively integrate the intrinsic properties of biological data into our KG-based model.

## RESULTS

To systematically verify the effectiveness of our method, a series of experiments are performed on the two above datasets. We first introduce the evaluation criteria and metrics. Then, the proposed method KGETCDA is compared with other SOTA methods. Moreover, ablation experiments are carried out to further evaluate the effectiveness of different modules. After that, we provide the best parameter settings of KGETCDA. Finally, we conducted case studies to evaluate the capability of our method for identifying potential associations between circRNAs and diseases. Through these experiments, we demonstrated the effectiveness of KGETCDA in accurately predicting potential CDA and capturing high-order interaction information with biological properties in an end-to-end manner.

### Evaluation criteria

Following previous studies (19, 28, 38), 5-fold cross-validation (CV) method is adopted to evaluate KGETCDA and compare with other SOTA models in this paper. Specially, in the 5-fold CV, all samples of the whole dataset are randomly divided into five equal parts and each part is selected for testing in turn while the remaining parts is utilized to train the model. Then, the prediction scores of test samples and unknown associations are obtained and sorted in descending order. The final results are the average values of five folds. Note that such a strategy helps reduce the over-fitting by fully using all samples in training. Finally, some classical evaluation metrics are applied to quantified the obtained results, including accuracy (*Acc*), precision (*Pre*), recall (*Rec*), F1-score (*F* 1). They are formulated as follows

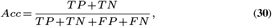

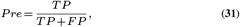

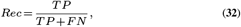

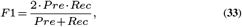

where *TP* and *TN* refer to the number of correctly predicted positive samples and negative samples, respectively; *FP* and *FN* refer to the number of incorrectly predicted positive samples and negative samples, respectively.

The area under the precision-recall curve (AUPR) and the area under the receiver operating characteristic curve (AUC) are also adopted to evaluate the method. AUPR is an effective metric for objectively indicating the changes of model performance and AUC is useful to represent model’s specificity and sensitivity, where the receiver operating characteristic curve (ROC) is plotted based on the true positive rate (*TPR*) and false positive rate (*FPR*) as follows:

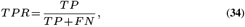

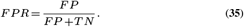

Moreover, existing methods usually use the number of correctly identified CDA to further measuring their prediction ability, which is formulated as Top-k.

### Performance comparison with SOTA methods

To demonstrate the superiority of our proposed method, we compare KGETCDA with 8 SOTAs, including network-based (KATZHCDA (11), RWR (14) and CD-LNLP (15)), traditional machine leaning-based (RNMFLP (19), DMFCDA (23)), and deep learning-based (GMNN2CD (27), KGANCDA (28), AE-RF (25)) methods. In the following part, we will compare them systematically from two perspectives, i.e. metrics comparison and statistical hypothesis test.

#### Metrics comparison across different datasets

With the same experimental settings, the final results of 5-fold CV on two datasets are obtained for nine models. The parameters of the other eight methods maintain consistency with their original paper. Note that these methods generally face test data leakage in their original implementation. In particular, information from the test set is used potentially during the model training of KGANCDA, GMNN2CD, RNMFLP and AE-RF, while GMNN2CD, RNMFLP, AE-RF obtain the similarity descriptor based on the whole dataset consisting of the train set and test set, and KGANCDA selects the best model with the help of model performance on test set. To address this problem, we create the similarity descriptor only based on the train set and only utilize the test set for testing.

For datset1, as illustrated in Figure 4A and 4B, KGETCDA consistently achieves the best performance in all metrics despite the sparsity of the dataset, which demonstrates our model’s outstanding prediction ability. The detailed information is listed in Supplementary Table S1, where KGETCDA obtains the best results of 0.4956 on *Acc*, 0.9228 on *Rec*, 0.0093 on *Pre*, 0.0173 *F* 1. Moreover, in order to further verify the prediction ability of models, the average number of correctly predicted CDA of five folds from top 10 to 40 is obtained, as shown in Figure 4C. Note that KGETCDA significantly yields the best performance (25.8, 35.8, 46, 54 for top 10 to top 40), which again demonstrates its strong prediction capability. More details are shown in Supplementary Table S2.

**Figure 4.**
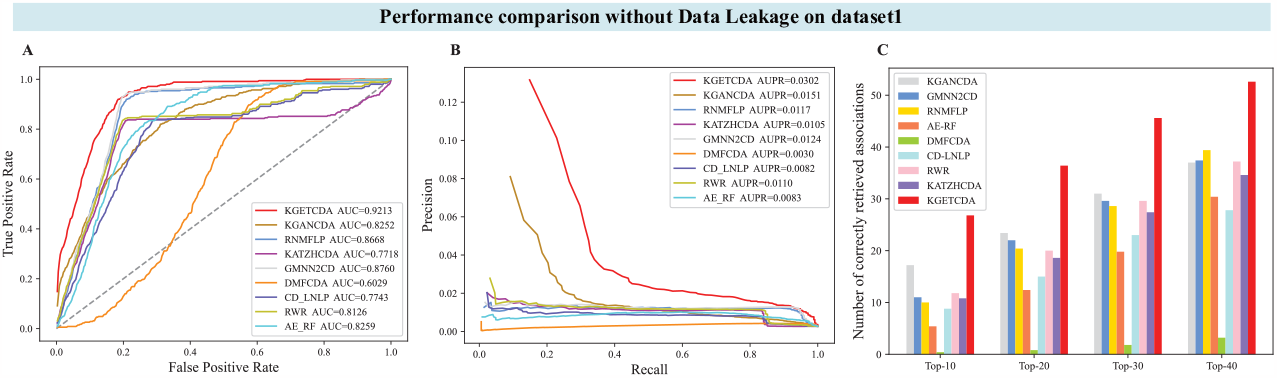
The comparison of our method with other predicting methods without data leakage problem on dataset1. **(A)** The AUC comparison **(B)** The AUPR comparison **(C)** The comparison of the number of correctly identified CDA.

For dataset2, as shown in Figure 5A and 5B, KGETCDA still obtains remarkable performance in AUC and AUPR, which again demonstrates our model’s strong prediction capability and excellent generalization. Additionally, we also take the average number of correctly predicted CDA of five folds from top 10 to 40 to further validate the prediction ability of our model on this large dataset. Noteworthy, as shown in 5C, our method significantly outperforms other methods (33.6, 51.2, 65.6, 78.4 for top 10 to top 40). On further analysis, due to the complex multi-relational scene, the performance of other methods is often limited. However, our method still maintains an outstanding prediction ability and produces reliable prediction results. More details are shown in the Supplementary File.

**Figure 5.**
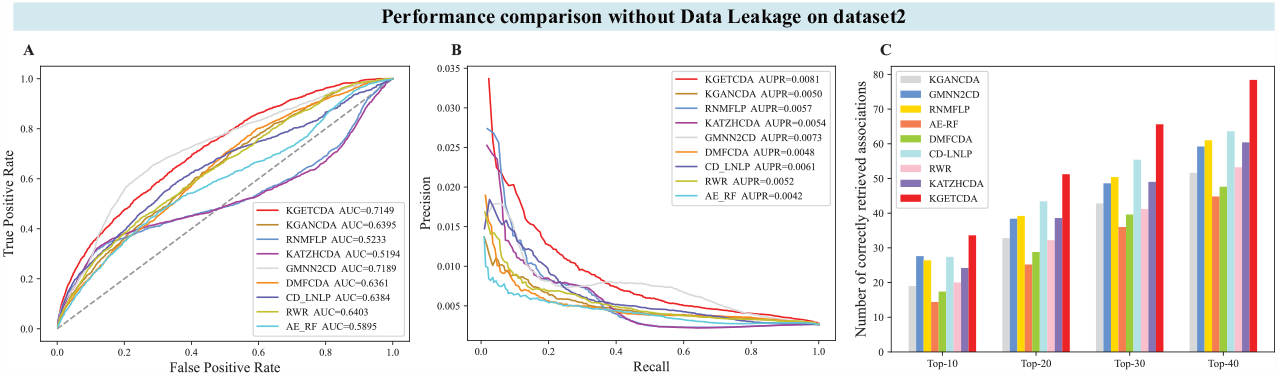
The comparison of our method with other predicting methods without data leakage problem on dataset2. **(A)** The AUC comparison **(B)** The AUPR comparison **(C)** The comparison of the number of correctly identified CDA.

In summary, our Transformer-based representation learning framework is more capable of identifying potential CDA than the baselines via accurately capturing high-order interaction information such as circRNA-lncRNA-disease, circRNA-lncRNA-miRNA-lncRNA-disease.

#### Statistical hypothesis test

To further evaluate the significant differences between KGETCDA and the baselines from statistical perspective, a nonparametric Wilcoxon signed-rank test-method (64) is applied, which is a powerful tool for making use of the magnitudes of the differences and thus effectively detect statistical differences between two models. The null hypothesis (*H*_0_) is considered that there is no obvious difference between KGETCDA and the baselines (*H*_0_ : *difference* = 0) while the alternative hypothesis (*H*_1_) is *H*_1_ : *difference* ≠ 0. Similar to (62, 63), we set the default P-value threshold to 0.05. As shown in Supplementary Table S3, all P-values between KGETCDA and other models are less than 0.05, and thus we reject the null hypothesis. Clearly, KGETCDA is superior to SOTA methods from a statistical perspective.

To summarize, the proposed method KGETCDA significantly outperforms all baselines from both biological and statistical perspectives. It’s our belief that this representation learning framework has a strong potential for effectively applying in other areas and lead to further improvement.

### Ablation Study

In this part, some ablation experiments are conducted to get deeper insights into the overall framework. We first study the performance of KGETCDA with different knowledge graph embedding (KGE) models and similarity-based negative sampling (NS) methods. Then, the effect of different tower structure of linear layers, dimensionality of embeddings, and the number of attention heads and layers in KRL are discussed. Finally, we explore the influence of different aggregators and scoring methods (illustrated in the Supplementary File). Here we present the abaltion results on dataset1, and results on dataset2 are available in Supplementary Figure S2.

#### Different KGE and similarity-based NS methods

To better verify the effectiveness of different modules, we conduct an ablation study on KGE and similarity-based NS modules. Specifically, we replace the Transformer-based KGE with commonly used baselines TransD (65), TransH (66) or ConvKB (67), named as TransD+INS, TransH+INS and ConvKB+INS, respectively (INS means integrated similarity-based NS). Moreover, different similairty-based NS methods are compared, where we replace NS part with GIPKS (named as Transformer+GNS), functional simialrity (Transformer+FNS), respectively. We obtain another variant by removing the similarity-based NS module and just use random NS strategy, named as Transformer+No NS. Note that only one module is replaced at once.

Five fold cross-validation AUC results are shown in Figure 6A. We observe that our design achieves slightly higher AUC than TransD+INS, TransH+INS and ConvKB+INS, which proves the powerful information extracting capability from multi-source data of our Transformer-based KRL. It’s promising that further exploring on this KRL framework in future work may lead to better improvements. Additionally, for the NS part, our design achieves remarkable performance, which indicates that integrating similarities from different representation spaces can effectively incorporate the intrinsic properties of biological data into KG and thus improve the representation learning. Note that all variants here outperform the current SOTA method GMNN2CD, which further demonstrate our method’s strong prediction capability.

**Figure 6.**
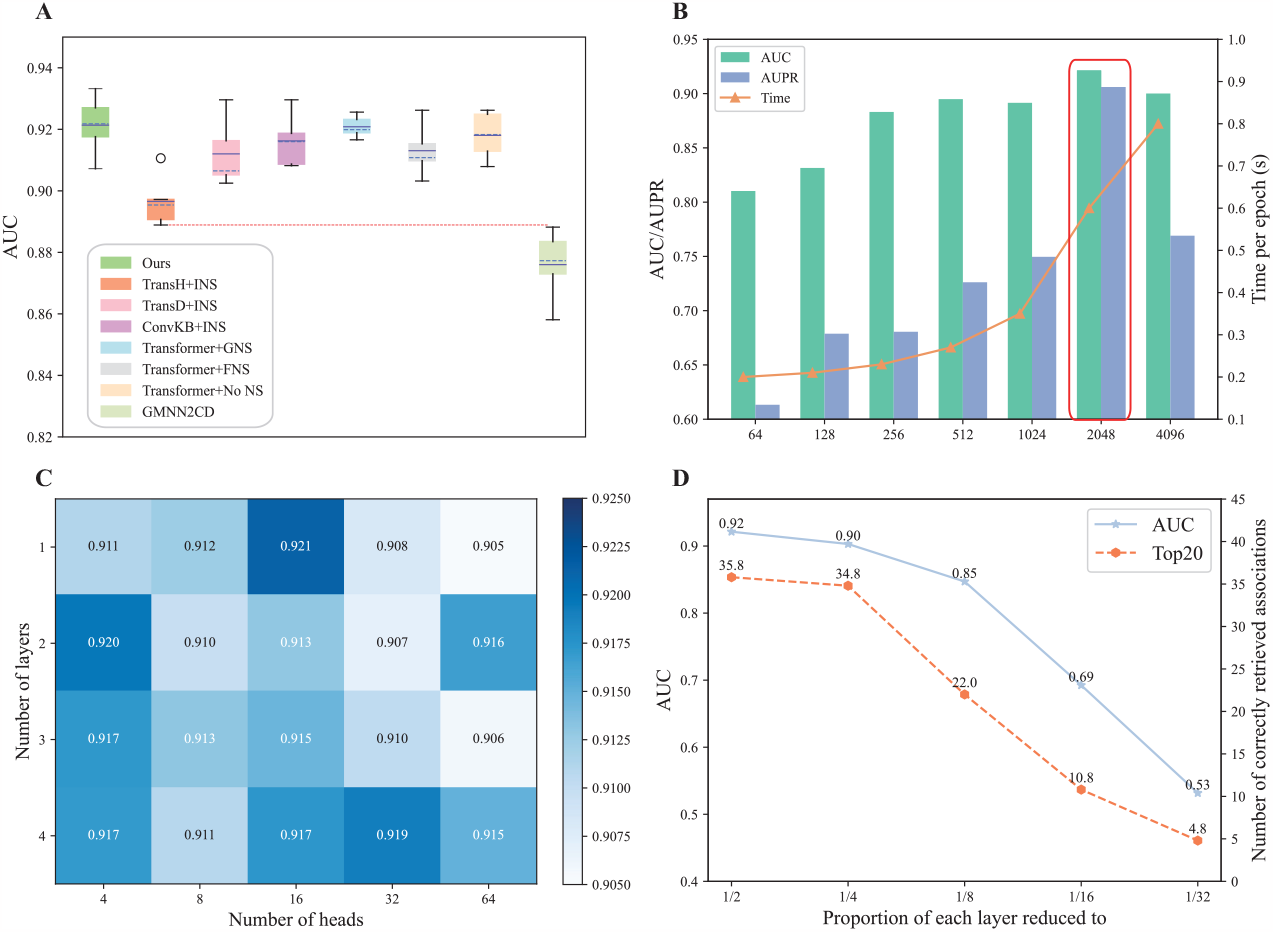
The results of ablation experiments on dataset1. **(A)** The effect of different knowledge graph embedding models and similarity-based negative sampling strategies. Our design achieves the best performance in both mean and median, and all variants here outperform the current SOTA method GMNN2CD. **(B)** The effect of different embedding dimensionality. We select 2048 here after considering both AUC and AUPR performance, and computational consumption. **(C)** The effect of the number of attention heads and layers in knowledge representation learning. **(D)** The difference of different tower structure of linear layers.

#### Effect of the embedding dimensionality

To explore the effect of the embedding dimensionality, we vary the size of dimensionality in 64, 128, 256, 512, 1024, 2048, 4096, which are termed with corresponding size (for example, size 64 is KGETCDA/64), and the results are illustrated in Figure 6B by scaling up the metric values for better visualization. Note that the performance gradually improves while increasing the size of embedding dimensionality until 4096. Specially, KGETCDA/2048 achieves the highest AUC and AUPR, which indicates that high embedding dimension enhance KGETCDA’s model learning capability and thus improve the performance. However, too high dimensions may introduce unusual features, lead to over-fitting problem and reduce generalization capabilities of the model.

#### Effect of the number of attention heads and layers in KRL

We further conduct a study to investigate the effect of the number of attention heads and layers in KRL. From Figure 6C we notice that KGETCDA achieves the best performance while the number of attention heads and layers are set to 16 and 1, respectively. while increasing the number attention heads from 4 to 16, the model performs better overall. It demonstrates that more attention heads facilitate more efficient representation learning and help encode more expressive information in different representation subspaces into embeddings than using multiple layers. Additionally, the performance of KGETCDA no longer increases and even degrades when attention heads increase from 16 to 64. This may be because large features lead to sparse feature representation and thus easily introducing useless information and even noise.

#### Tower structure of linear layers

As illustrated in (20, 61), reducing the number of hidden units after each hidden layer can facilitate more abstractive features learning and the tower structure has been proven to be particularly efficient and suitable, which reduces 1*/*2^*n*^ to for each layer. Specially, we consider that the reduction for each layer is searching in 1/2, 1/4, 1/8, 1/16, 1/32. From Figure 6D, we observe that KGETCDA with halving the layer size is superior to others and we use this structure in our research.

### Parameter setting

We implement KGETCDA in PyTorch with an NVIDIA Titan Xp GPU and optimize the whole model with Adam (60) optimizer. The learning rate and weight decay on all parameters are set to 1e-4 and 1e-5 with 100 training epochs. To accurately exploit high-order features and obtain high-quality embeddings, the dimension of each propagation layer is set to 512, 256, 128, 64, respectively. To reduce the over-fitting on the train set, we apply dropout after each propagation layer with the same rate of 0.1. We also apply dropout to the input of KRL, the output of scaled dot product, multi-head attention and feed-forward network with a rate of 0.2, 0.3, 0.2, 0.3 by default, respectively. Additionally, after extensive experiments, our method achieves the best performance while the ratio of negative to positive samples is set to 8:1 and the number of similar-based sampling is 10. The detailed information of parameter setting are listed in Supplementary Table S6.

### Case studies

To further evaluate the effectiveness of KGETCDA, case studies are conducted on both two datasets. Following (23, 38, 62), we train the model KGETCDA on all known CDA, and then predict potential probabilities for all unknown CDA and sort them in descending order. Moreover, we collect experimental evidence to verify them in public databases and newly published literatures, and the results are listed in Table 2.

**Table 2.** The top-10 predicted lung cancer or leukemia related candidate circRNAs

| Dataset | Disease | Candidate circRNA | Rank | Evidence (PMID) |
| --- | --- | --- | --- | --- |
| Dataset1 | Leukemia | hsa_circ_0075001 | 1 | 28971903 |
|  |  | circpan3 | 2 | 31401408, 30395908 |
|  |  | circ-anapc7 | 3 | 29969755 |
|  |  | hsa_circ_100290 | 4 | unknown |
|  |  | hsa_circ_0035381 | 5 | 28282919 |
|  |  | circznf609 | 6 | 33259601 |
|  |  | hsa_circ_0017446 | 7 | 28282919 |
|  |  | hsa_circ_0000488 | 8 | 30037980 |
|  |  | hsa_circ_0049657 | 9 | 28282919 |
|  |  | hsa_circ_0008078 | 10 | 28282919 |
| Dataset2 | Lung Cancer | CDR1as | 1 | 30022841 |
|  |  | hsa_circ_0007534 | 2 | 30017736 |
|  |  | circSOX4 | 3 | 32112500 |
|  |  | circWHSC1 | 4 | 34551195, 29722168 |
|  |  | hsa_circ_0000735 | 5 | 30737027 |
|  |  | hsa_circ_0015449 | 6 | unknown |
|  |  | hsa_circ_0075829 | 7 | unknown |
|  |  | hsa_circ_0067934 | 8 | 29863250, 30344708 |
|  |  | circSMARCA5 | 9 | 33085761 |
|  |  | hsa_circ_0025033 | 10 | unknown |

For dataset1, we choose leukemia, which has affected human health severely, and existing studies have demonstrated that the studies of circRNA benefit the treatment of leukemia. From Table 2, we can observe that top-10 leukemia-related leukemia, and 9 of 10 circRNAs (hsa circ 0075001, circpan3, circ-anapc7, hsa circ 0035381, circznf609, hsa circ 0017446, hsa circ 0000488, hsa circ 0049657, hsa circ 0008078) are confirmed. For example, circpan3 mediates drug resistance in leukemia via the miR-183-5p-XIAP/miR-153-5p axis (68).

Hsa circ 0000488 accelerates human leukemia progression via promoting PRKACB expression and suppressing microRNA 496 (69).

For dataset2, we choose lung cancer (LC), which is a common cancer worldwide, and accurately identifying breast cancer-related circRNAs can offer new insights into disease diagnosis and treatment. As shown in Table 2, top-10 lung cancer-related circRNAs are selected based on their predicted scores, and 7 of 10 circRNAs (CDR1as, hsa circ 0007534, circSOX4, circWHSC1, hsa circ 0000735, hsa circ 0067934, circSMARCA5) are confirmed by existing literatures. For example, as an oncogene, overexpressed CDR1as ranked at top 1 can promote the tumor progression via miR-7 in lung cancer (70). The expression level of hsa circ 0067934 ranked at top 8 affects the proliferation and metastasis of LC (71).

In summary, case studies conducted on two datasets demonstrate our proposed method’s significant role in identifying unknown CDA, and it deserves experimental tests to further verify these CDA.

## WEB-BASED VISUALIZATION

Considering that existing computational methods often present results and offer collected data resources only in the form of static figures or tables, which may not be user-friendly relatively for researchers to use, we develop an online interaction web-based platform named HNRBase, which provides intuitive visualization and user-friendly web interaction interfaces for intelligent search, convenient data download, and reliable model prediction. In particular, as illustrated in Figure 7, our web interaction interface mainly consists of 4 core functions: intelligent search and browse, model prediction, information visualization, and advanced interaction. The detailed architecture design and description are illustrated in the Supplementary File. Note that our web-based platform enables novel visualization, accessible resources and user-friendly interaction. This design enhances user experience well and benefits further research in the field of CDA prediction. Moreover, to further utilize our method, we expand this framework to miRNA-disease and lncRNA-disease prediction due to the high similarity between them and CDA prediction.

**Figure 7.**
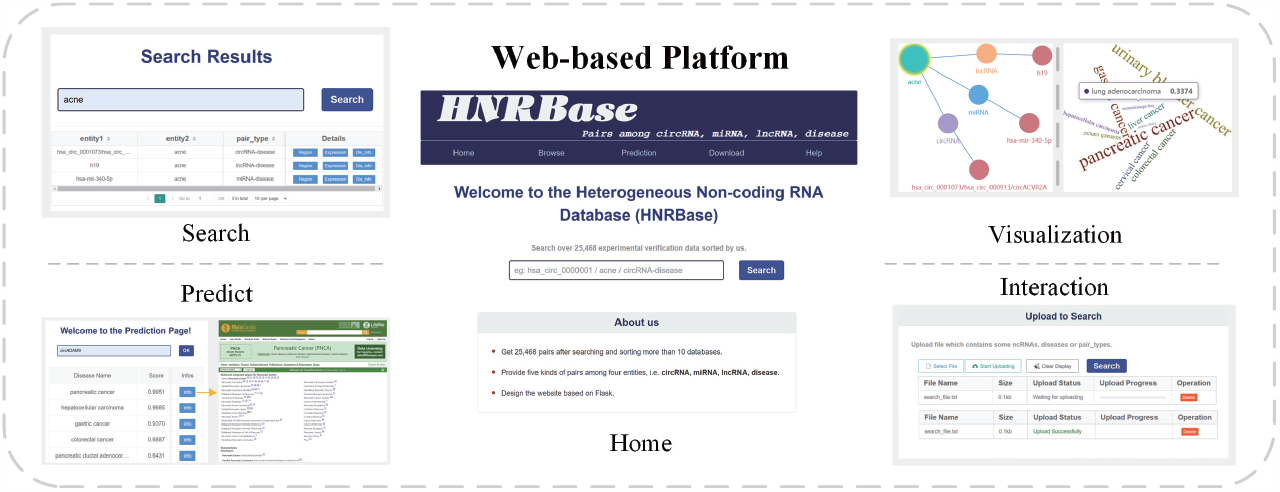
The overview of web-based platform.

## DISCUSSION

Increasing studies have demonstrated that circRNA plays a significant role in the initiation and progression of complex human diseases, exhibiting rich biological functions. Effectively predicting potential CDA can offer novel insights into disease diagnosis and treatment. However, existing computational methods face challenges in effectively exploring high-order interaction information that incorporates biological intrinsic properties due to data sparsity problem. In this paper, we propose KGETCDA, a new CDA prediction method based on a Transformer-based knowledge graph representation learning framework, which produces high-quality feature representations. Empirical experiments conducted on two heterogeneous datasets reveal that KGETCDA significantly outperforms state-of-the-art (SOTA) methods, and accurately captures high-order information with biological properties in an end-to-end manner. Moreover, we develop an online interaction web-based platform named HNRBase, which provides user-friendly interfaces for intuitive visualization, intelligent search, convenient data download, and reliable model prediction. This presentation manner of results enhances user experience well and benefits further research in the field of CDA prediction.

Furthermore, as there are still some promising avenues for further research, we suggest several intuitive improvements worth exploring. Firstly, more biological data can be integrated to further verify KGETCDA’s effectiveness, and we hope that this representation learning framework can open up future research probabilities for other areas such as drug-protein prediction and gene-RNA prediction.

Secondly, more efficient scoring methods in Transformer-based knowledge representation learning can be introduced to further improve performance. Thirdly, incorporating additional biological auxiliary knowledge, such as RNA sequence and gene expression information, into KG would be a promising approach. Finally, due to Transformer’s intensive computational and memory consumption, we anticipate investing more efforts to tackle this problem in the future.

## Supporting information

This is the supplementary file of KGETCDA.

## DATA AVAILABLE

The code and datasets are publicly available at https://github.com/jinyangwu/KGETCDA.

## SUPPLEMENTARY DATA

Supplementary data are available online at https://academic.oup.com/nar.

## AUTHOR CONTRIBUTIONS STATEMENT

J.W. and L.F. conceived the project, conducted the experiments, analysed the results, and wrote the manuscript. Z.N., Y.D., Y.W. and Q.P. reviewed the manuscript.

## ACKNOWLEDGEMENTS

We acknowledge helpful discussions with members of the Peng lab.

