## Supplementary material for "KGETCDA: an efficient representation learning framework based on knowledge graph encoder from transformer for predicting circRNA-disease associations": This is the supplementary file of KGETCDA.

### Appendix

### Supplementary Information of Methods

#### 1 Performance comparison

In this part, we will introduce some experimental details. As shown in Table S1 and Table S2, we list the detailed information of performance comparison of nine models, where KGETCDA consistently achieves the best performance on two datasets. Moreover, the statistical significance of differences between KGETCDA and the other eight methods is shown in Table S3, which further demonstrates our method's efficiency in accurately identifying potential circRNA-disease associations.

#### 2 Supplementary ablation experiments

For the ablation part, the results of dataset2 are shown in Figure S2. Moreover, to verify the effectiveness of GCN aggregator in KGETCDA, we conduct an ablation study by replacing it with GraphSage or Bi-Interaction. As shown in Table S4, KGETCDA with GCN performs better than GraphSage and Bi-Interaction aggregator overall. One possible reason is that GCN aggregator fully exploit the interaction information between an entity and its ego-network, while Bi-Interaction introduces noise and GraphSage ignores some interaction information.

Additionally, the results of different scoring functions are shown in Table S5. Here we compare TwoMult ( $f = e_r e_t$ ), ThreeMult ( $f = e_h e_r e_t$ ), and Distance-based ( $f = e_h + e_r - e_t$ ) scoring methods. Given that TwoMult performs slightly better than ThreeMult and Distance-based scoring methods overall, we select the scoring function TwoMult in our research.

#### 3 Web-based platform

As shown in Figure S3, Our web interaction interface mainly consists of 4 core functions: intelligent search and browse, model prediction, information visualization, and advanced interaction. The homepage is shown in Figure S4 and the detailed description is illustrated as follows.

##### *3.1 Intelligent search and browse, and user-friendly visualization*

As shown in Figure S5, the 'Search' function allows users to efficiently retrieve 4 entities (circRNA, miRNA, lncRNA, disease), and 5 relations (circRNA-disease, miRNA-disease, lncRNA-disease, circRNA-miRNA, miRNA-lncRNA) via inputting their names or ids and clicking the 'Search' button. The requests will be submitted to the backend server and then corresponding search results are displayed in the form of both tables and multi-relational graphs, which effectively reflects the complicated relations of the input key words. Note that users can reposition nodes in graphs on-the-fly by clicking and dragging and they will be highlighted while hovering over them, which enhance user experience well. Additionally, according to searching entries, our platform is also capable of automatically searching and matching with UCSC Genome Browser, EMBL-EBI and MalaCards databases to visualize the gene location information, the profiles of upstream and downstream regulatory elements, the corresponding gene expression and disease information, which incorporates more useful information into our platform for researchers working in different fields to use.

As shown in Figure S5, users can also easily browse all entities and relations through clicking the tab 'Browse'. Then, an interactive wordcloud diagram of part data based on the degree of the entities in the multi-relational graphs is obtained, which provides a more comprehensive presentation of the overall database.

##### *3.2 Model prediction and information visualization*

The 'model prediction' module refers to the prediction of associations between circRNAs and diseases that have not been formally documented or experimentally validated. Through deploying the proposed prediction model KGETCDA, potential unknown CDAs are predicted and presented in the form of tables as well as interactive wordcloud diagrams. Specifically, users can simply input circRNAs of interest and click the 'OK' button to the backend cloud server, and then the top-10 prediction results will be quickly displayed in a table. The detailed information of corresponding diseases is also available through clicking the 'detail' button. Furthermore, a wordcloud diagram is presented beside the table and the predicted probability of each CDA is displayed while hovering over candidate diseases (Figure S6). Furthermore, we also expand our model to predict miRNA-disease and lncRNA-disease due to the high similarity between them and circRNA-disease prediction.

##### *3.3 Advanced interaction*

To further satisfy user's requirements, we design some advanced interaction functions, including file upload and download, data transfer, and visual interaction functions. As illustrated in Figure S5 and Figure S7, our platform supports users in uploading files of interest to obtain relevant search entries and prediction results, which are available for download. We also provide a user-friendly interface for convenient download of the integrated dataset collected before for further research in the field of CDA prediction. Finally, users can easily experience visual interaction in the 'Search' and 'Prediction' webpage.

### Tables

**Table S1**. The detailed information of the results of nine models on two datasets

| Dataset | Methods | Acc | Pre | Recall | F1 |
|---|---|---|---|---|---|
| Dataset1 | **KGETCDA** | **0.4956** | **0.9228** | **0.0093** | **0.0173** |
| | KGANCDA (2022) | 0.4950 | 0.8278 | 0.0071 | 0.0134 |
| | GMNN2CD (2022) | 0.4953 | 0.8779 | 0.0068 | 0.0132 |
| | RNMFLP (2022) | 0.4953 | 0.8687 | 0.0066 | 0.0129 |
| | AE-RF (2021) | 0.4950 | 0.8284 | 0.0057 | 0.0111 |
| | DMFCDA (2021) | 0.4938 | 0.6084 | 0.0028 | 0.0056 |
| | CD-LNLP (2019) | 0.4948 | 0.7774 | 0.0056 | 0.0109 |
| | RWR (2018) | 0.4950 | 0.8153 | 0.0064 | 0.0124 |
| | KATZHCDA (2018) | 0.4947 | 0.7748 | 0.0061 | 0.0119 |
| Dataset2 | **KGETCDA** | **0.4953** | 0.7182 | **0.0052** | **0.01** |
| | KGANCDA (2022) | 0.4949 | 0.6438 | 0.0041 | 0.008 |
| | GMNN2CD (2022) | 0.4953 | 0.7222 | 0.0051 | 0.0098 |
| | RNMFLP (2022) | 0.4943 | 0.5288 | 0.0039 | 0.0076 |
| | AE-RF (2021) | 0.4946 | 0.5944 | 0.0037 | 0.0073 |
| | DMFCDA (2021) | 0.4949 | 0.6404 | 0.0037 | 0.0073 |
| | CD-LNLP (2019) | 0.4949 | 0.6425 | 0.0045 | 0.0086 |
| | RWR (2018) | 0.4949 | 0.6446 | 0.0042 | 0.0082 |
| | KATZHCDA (2018) | 0.4943 | 0.5249 | 0.0039 | 0.0075 |

**Table S2**. The number of correctly identified associations of all models on two datasets

| Dataset | Methods | TOP-10 | TOP-20 | TOP-30 | TOP-40 |
|---|---|---|---|---|---|
| Dataset1 | **KGETCDA** | **25.8** | **35.8** | **46** | **54** |
| | KGANCDA (2022) | 17.2 | 23.4 | 31 | 37 |
| | GMNN2CD (2022) | 11 | 22 | 29.6 | 37.4 |
| | RNMFLP (2022) | 10 | 20.4 | 28.6 | 39.4 |
| | AE-RF (2021) | 5.4 | 12.4 | 19.8 | 30.4 |
| | DMFCDA (2021) | 0.4 | 0.8 | 1.8 | 3.2 |
| | CD-LNLP (2019) | 8.8 | 15 | 23 | 27.8 |
| | RWR (2018) | 11.8 | 20 | 29.6 | 37.2 |
| | KATZHCDA (2018) | 10.8 | 18.6 | 27.4 | 34.6 |
| Dataset2 | **KGETCDA** | **33.6** | **51.2** | **65.6** | **78.4** |
| | KGANCDA (2022) | 19 | 32.8 | 42.8 | 51.6 |
| | GMNN2CD (2022) | 27.6 | 38.4 | 48.6 | 59.2 |
| | RNMFLP (2022) | 26.4 | 39.2 | 50.4 | 61 |
| | AE-RF (2021) | 14.4 | 25.2 | 36 | 44.7 |
| | DMFCDA (2021) | 17.4 | 28.8 | 39.6 | 47.6 |
| | CD-LNLP (2019) | 27.4 | 43.4 | 55.4 | 63.6 |
| | RWR (2018) | 20 | 32.2 | 41.2 | 53.2 |
| | KATZHCDA (2018) | 24.2 | 38.6 | 49 | 60.4 |

**Table S3**. Statistical significance of differences between KGETCDA and the other eight methods

| | Dataset | KGANCDA | GMNN2CD | RNMFLP | AE-RF | DMFCDA | CD-LNLP | RWR | KATZHCDA |
|---|---|---|---|---|---|---|---|---|---|
| P-value | Dataset1 | 1.102e-10 | 1.103e-10 | 1.104e-10 | 1.107e-10 | 1.105e-10 | 1.102e-10 | 1.097e-10 | 1.099e-10 |
| | Dataset2 | 1.105e-10 | 1.894e-6 | 1.104e-10 | 1.106e-10 | 1.624e-10 | 1.104e-10 | 1.105e-10 | 1.106e-10 |

**Table S4**. Effect of aggregators

| Dataset | Aggregator | AUC | AUPR | Top-10 |
|---|---|---|---|---|
| Dataset1 | **GCN** | 0.9213 | **0.0302** | 25.8 |
| | GraphSage | 0.9134 | 0.0252 | 22.6 |
| | Bi-Interaction | 0.9228 | 0.0274 | 26 |
| Dataset2 | **GCN** | **0.7149** | **0.0081** | **33.6** |
| | GraphSage | 0.6957 | 0.0078 | 32 |
| | Bi-Interaction | 0.712 | 0.0077 | 30.2 |

**Table S5**. Effect of scoring functions

| Dataset | Aggregator | AUC | AUPR | Top-10 |
|---|---|---|---|---|
| Dataset1 | **TwoMult** | **0.9213** | **0.0302** | 25.8 |
| | ThreeMult | 0.9213 | 0.0288 | 26.2 |
| | Distance-based | 0.9204 | 0.0289 | 26.0 |
| Dataset2 | **TwoMult** | 0.7149 | **0.0081** | **33.6** |
| | ThreeMult | 0.7151 | 0.0079 | 31 |
| | Distance-based | 0.7144 | 0.0079 | 32.2 |

**Table S6**. Summaries of experimental settings

| Hyperparameter | Our settings |
|---|---|
| Training epochs | 100 |
| Lr | 0.0001 |
| Batch size | 256 |
| Dim features | 2048 |
| Dim of hidden layer in FFN | 512 |
| Transformer-based encoder layers | 1 |
| Attention heads | 16 |
| Aggregation layers & Dim | 4, [512, 256, 128, 64] |
| Weight decay | 1e-5 |
| Knowledge graph dropout | [0.2, 0.3, 0.2, 0.3] |
| Aggregation layer dropout | [0.1, 0.1, 0.1, 0.1] |
| MLP dropout | 0.04 |
| The ratio of negative sampling | 8:1 |
| K for top-k similar circRNAs in negative sampling | 10 |

### Figures

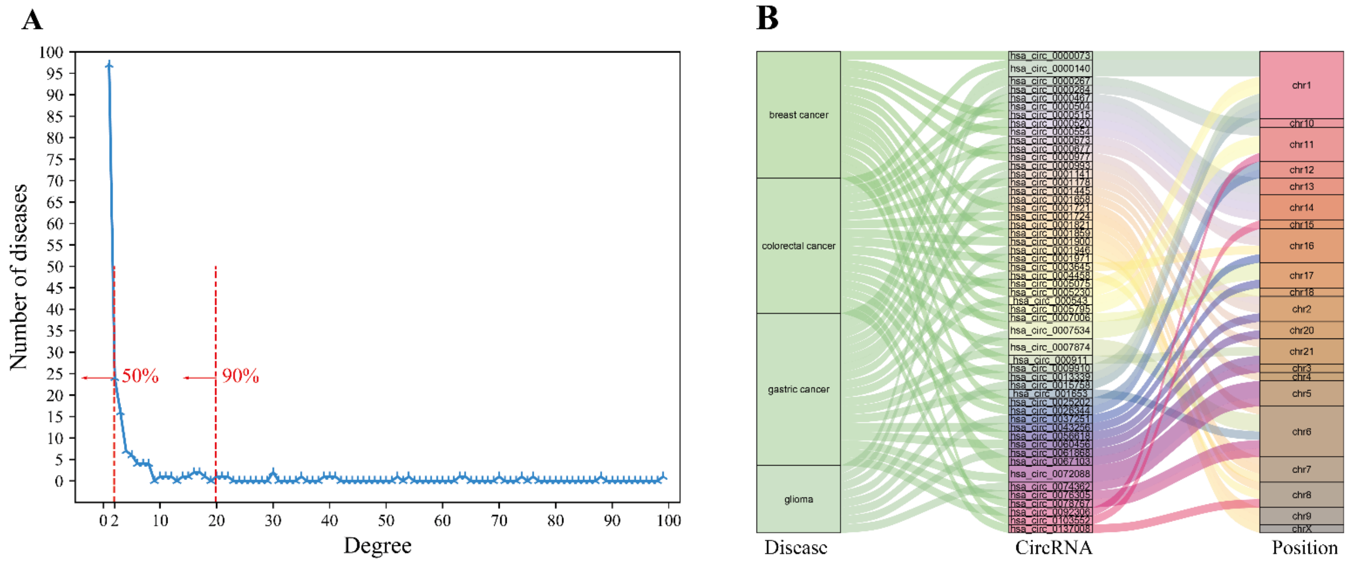

**Figure S1.** Visualization of dataset2. **(A)** The degree distribution. It represents the number of corresponding diseases associating with different numbers of circRNAs in dataset2. **(B)** The visualization of sankey diagram of some associations in dataset2.

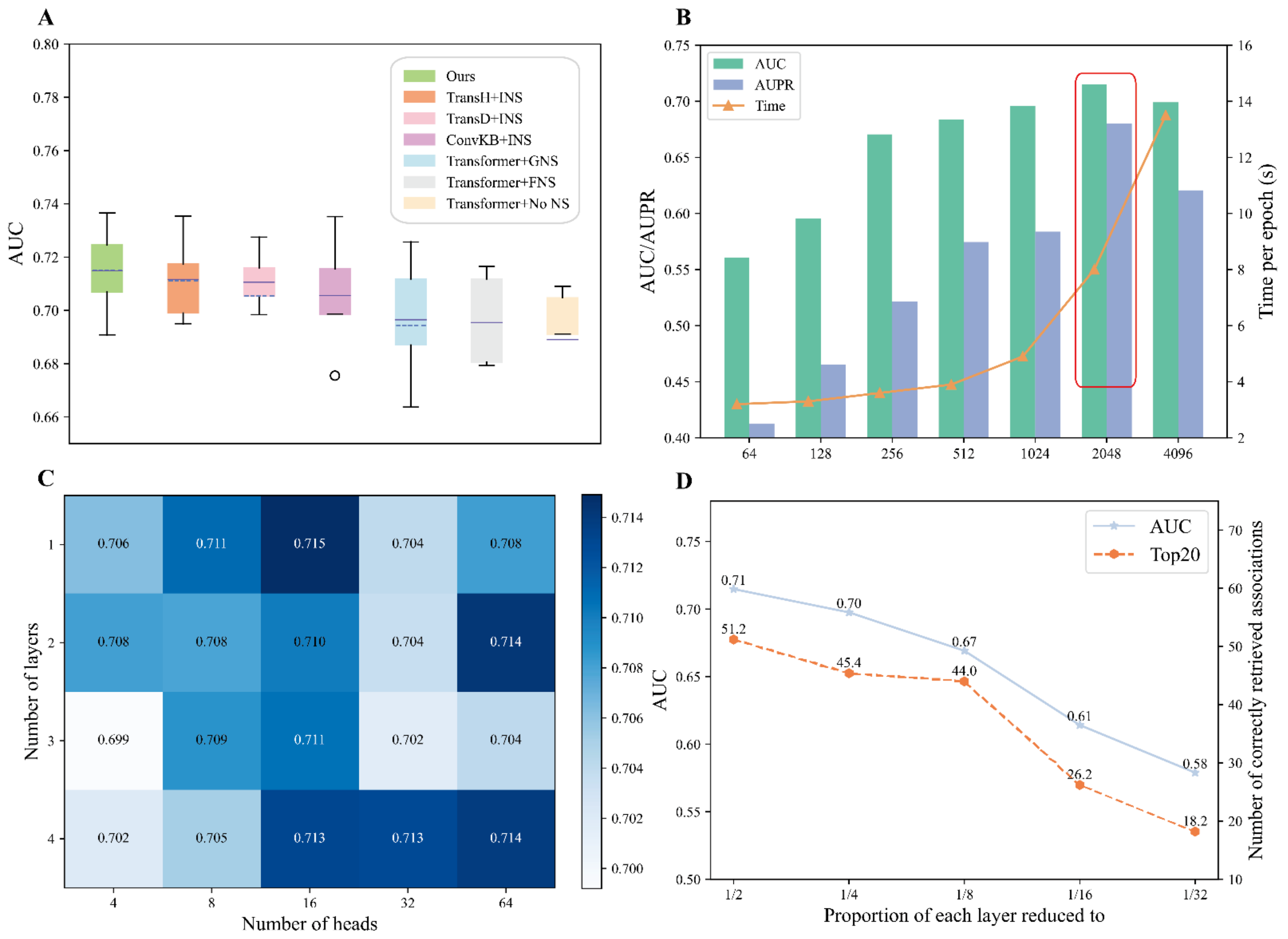

**Figure S2.** The results of ablation experiments on dataset2. **(A)** The effect of different knowledge graph embedding models and similarity-based negative sampling strategies. Our design achieves the best performance in both mean and median. **(B)** The effect of different embedding dimensionality. We select 2048 here after considering both AUC and AUPR performance, and computational consumption. **(C)** The effect of the number of attention heads and layers in knowledge representation learning. **(D)** The difference of different tower structure of linear layers.

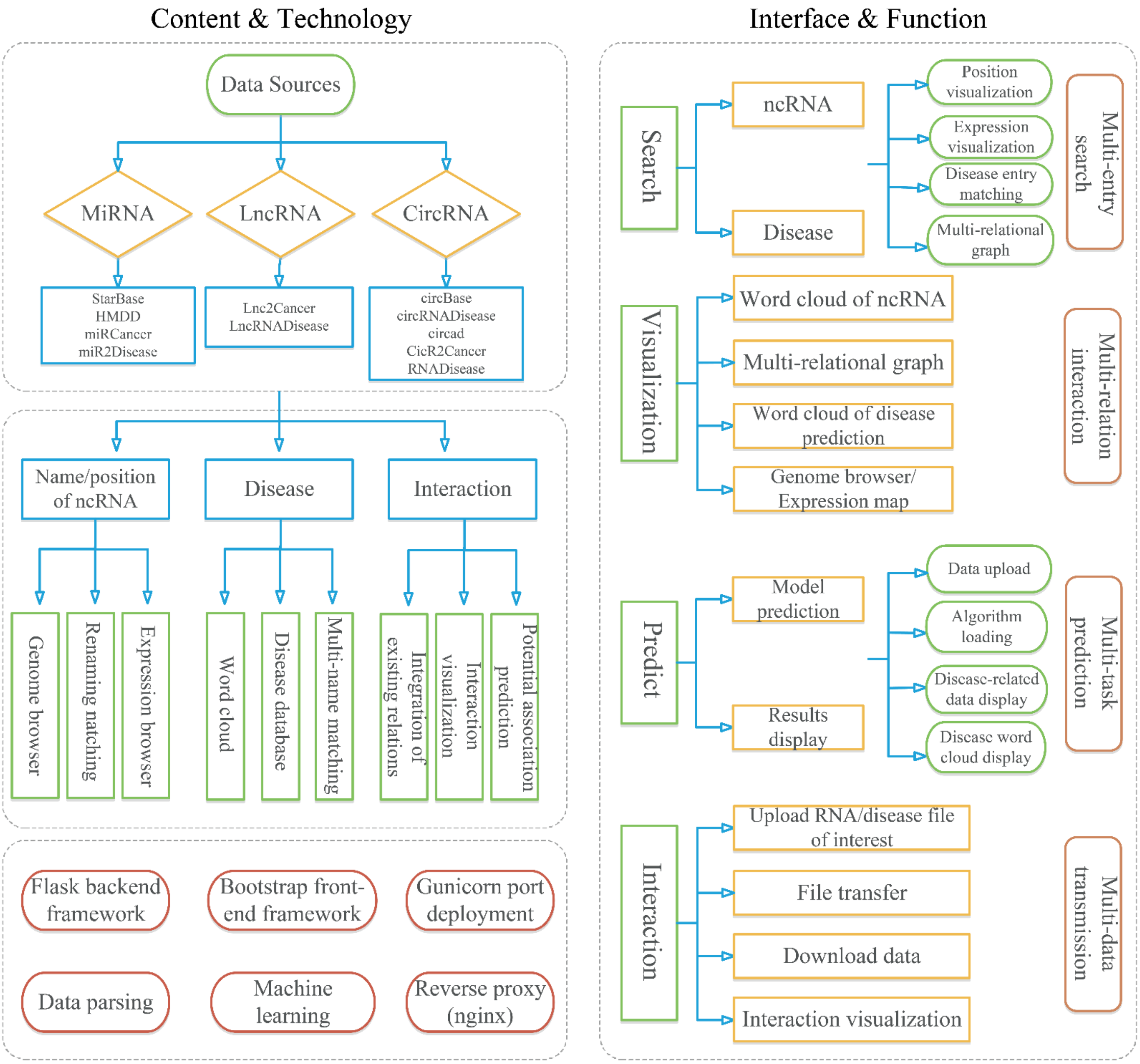

**Figure S3.** The framework of our web-based platform, which contains 4 parts: search, predict, visualization, interaction.

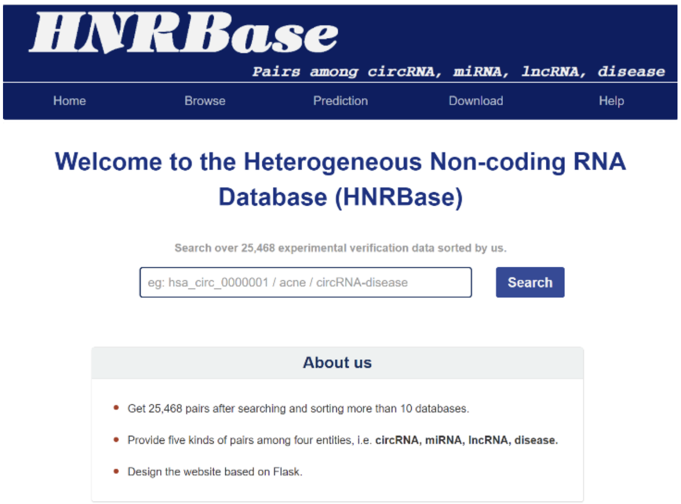

**Figure S4.** The homepage of our web-based platform, where illustrates the basic information of our platform, such as the total number of collected data and related databases.

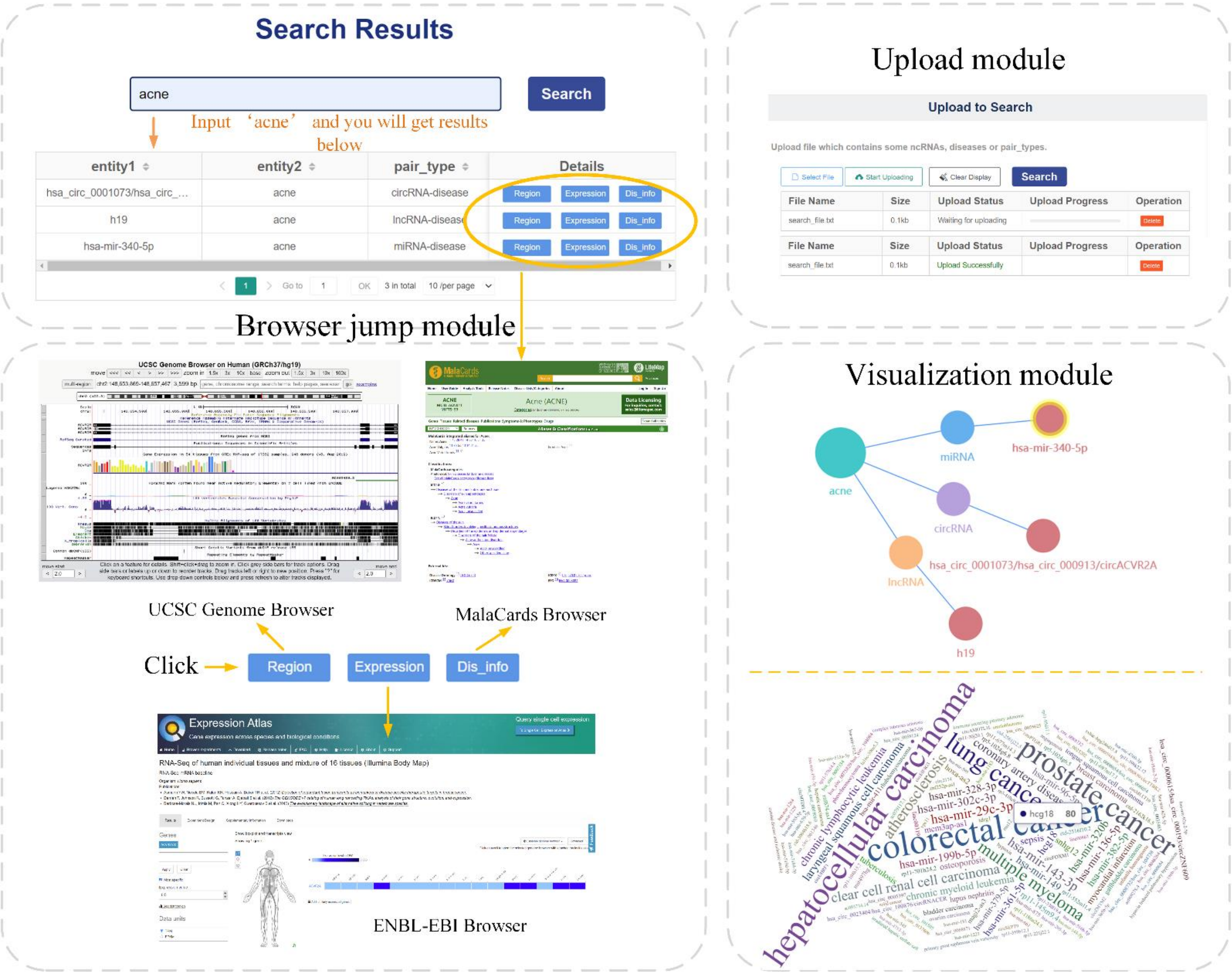

**Figure S5.** The search and browse module. Input a candidate key and you will get results in the form of tables and interactive multi-relational graphs. Moreover, you can also fully use the browser jump module and get further information. An interactive wordcloud diagram is also shown in the browser module.

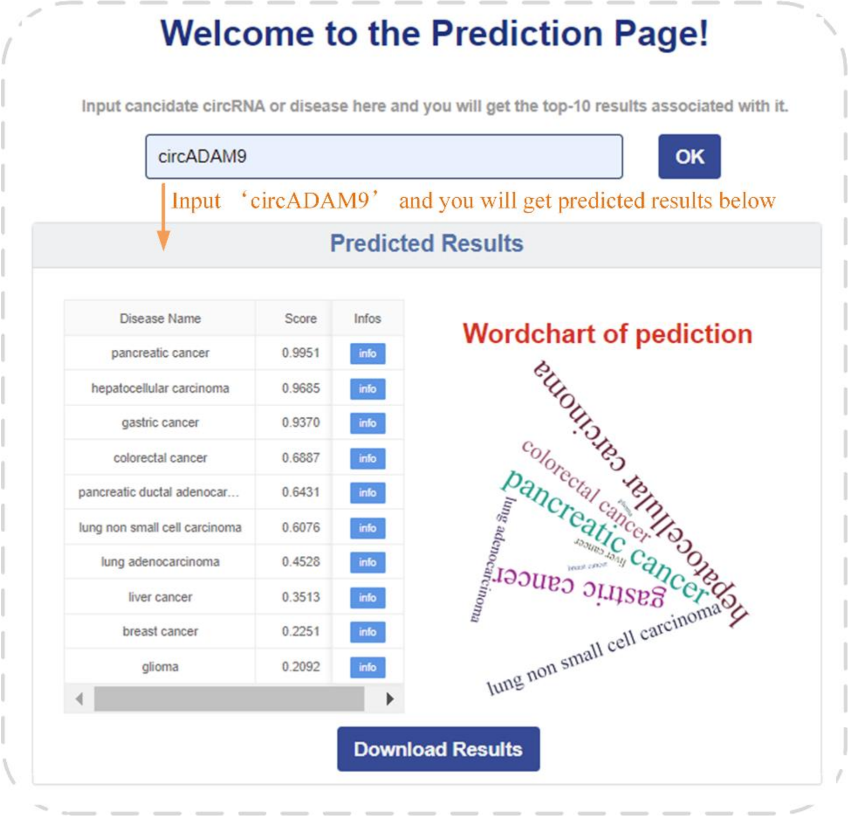

**Figure S6.** The prediction module. Input a candidate key and you will get prediction results in the form of tables and interactive wordcloud diagrams. Moreover, you can also upload file to obtain batch results at once.

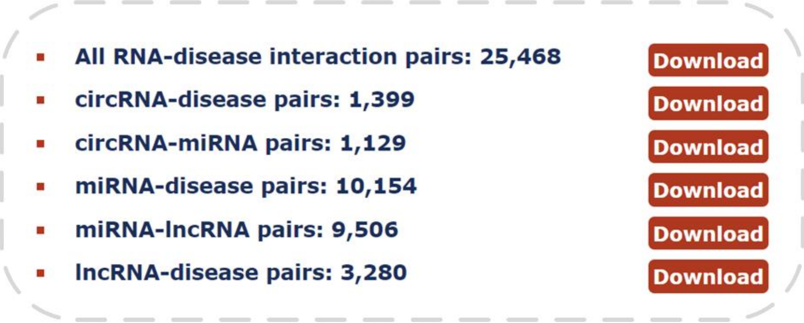

**Figure S7.** The download module. You can get results of interested entities in the form of CSV file. We provide multiple forms of download.
